# Dietary Fat Content Influences PanIN Progression and Pancreatic Cancer Development in Mice

**DOI:** 10.1101/2025.11.26.690783

**Authors:** Urvinder Kaur Sardarni, Erika Y. Faraoni, Alyssa M. Waller, Lincoln N. Strickland, Baylee O’Brien, Jesse L. Cox, Florencia McAllister, Jennifer M. Bailey-Lundberg

**Affiliations:** Department of Pathology, Microbiology and Immunology, University of Nebraska Medical Center, Omaha, NE, USA; Fred and Pamela Buffett Cancer Center, University of Nebraska Medical Center, Omaha, NE, USA; Department of Genetics, University of Texas MDAnderson Cancer Center, Houston, TX, USA; The Eppley Institute for Research in Cancer and Allied Diseases, University of Nebraska Medical Center, Omaha, NE, USA; Texas A&M Medical School, College Station, TX, USA; GI Medical Oncology, The University of Texas MD Anderson Cancer Center, Houston, TX, USA

## Abstract

Dietary macronutrient composition has emerged as a key modulator of pancreatic tumorigenesis, yet the impact of lipid-rich diets, particularly ketogenic diets (KD) on the earliest stages of pancreatic cancer development remains unclear. To investigate how dietary lipids shape the initiation and progression of Kras-driven neoplasia, we examined the effects of low-fat diet (LFD), high-fat diet (HFD), and KD in the *Ptf1a^CreERT2^;Kras^G12V^ (Acinar^KrasG12V^)* mouse model. KD-fed mice showed the shortest survival (median 26 ± 7 days) compared with SD (87 ± 29; p = 0.02) and LFD (57 ± 27; p = 0.02), while HFD-fed mice also exhibited reduced survival relative to SD (35 ± 25; p = 0.05). KD feeding induced severe glucose intolerance and elevated circulating β-hydroxybutyrate levels. Histologically, KD-fed *Acinar^KrasG12V^* mice developed invasive, sarcomatoid-like pancreatic ductal adenocarcinoma (PDAC), while HFD-fed mice showed increased poorly differentiated PDAC; in both groups these aggressive tumors were associated with extensive fibrosis and increased stromal CD39 expression relative to tumor compartments. Proteomic analysis demonstrated activation of PI3K–Akt–mTOR and EGFR signaling in KD and HFD-fed *Acinar^KrasG12V^* mice. Serum cytokines/chemokines profiling revealed pro-inflammatory and pro-angiogenic milieu in KD-fed *Acinar^KrasG12V^*mice. Collectively, these results show that dietary lipid enrichment prior to oncogenic Kras activation may accelerate early pancreatic neoplasia and foster a microenvironment conducive to tumor progression. These findings underscore the need for careful consideration of KD use in individuals at elevated risk for pancreatic cancer.

## INTRODUCTION

Dietary macronutrient composition has emerged as a key modifier of pancreatic tumorigenesis. In *Kras^G12D^* mice, high-fat diet (HFD) feeding accelerates pancreatic intraepithelial neoplasia (PanIN) formation, enhances inflammation and desmoplasia, and reduces survival (1, 2). High-carbohydrate diets (HCDs) exert similar but less pronounced effects, whereas high-protein diets (HPDs) show negligible impact on pancreatic neoplasia compared with controls (3). Interestingly, ketogenic diets (KDs), composed of 80–90% fat, have demonstrated anti-tumor effects across several tumor types in both clinical and preclinical studies (4). By elevating circulating ketone bodies, KDs reprogram cellular metabolism toward fatty acid and ketone utilization, while limiting glucose availability. Since cancer cells rely heavily on aerobic glycolysis to sustain growth, restricting carbohydrates has been proposed as a metabolic strategy to suppress tumor progression (5). However, the impact of KD in PDAC appears context dependent. When combined with standard chemotherapies or administered to obese *Kras^G12D^*mice, KD can improve survival and delay tumor progression (6–8). In contrast, KD accelerated tumor growth in lean *Kras^G12D^*models (7). These findings underscore the complexity of diet–tumor interactions and highlight that the preventive potential of KD in pancreatic cancer remains uncertain.

Recent transcriptomic analyses of healthy donor pancreata have revealed the unexpected presence of PanIN lesions, suggesting that early neoplastic lesions may be common and potentially amenable to dietary modulation (9). To explore how dietary lipid content influences early pancreatic tumorigenesis, we investigated the effects of KD, HFD, and low-fat diet (LFD) feeding on pancreatic disease progression using the *Ptf1a^CreERT2^;Kras^G12V^ (Acinar^KrasG12V^)* mouse model, which allows temporal control of oncogenic Kras activation in acinar cells. Mice were preconditioned with distinct dietary regimens prior to Kras activation, and were monitored for survival, histopathological progression, and inflammatory and molecular signaling.

Here, we demonstrate that dietary composition prior to oncogenic Kras activation profoundly shapes pancreatic cancer trajectory. While HFD predictably accelerates disease, KD unexpectedly promotes more aggressive pathology and reduced survival compared with LFD and standard chow diet (SD). Mechanistically, KD and HFD feeding activated PI3K–Akt–mTOR and EGFR signaling pathways and induced a systemic pro-inflammatory, pro-angiogenic cytokine milieu. These findings reveal how dietary lipids can potentiate oncogenic and inflammatory pathways in the pancreas, offering new insight into the dietary modulation of pancreatic cancer risk.

## RESULTS

### KD and HFD reduce survival in *Acinar^KrasG12V^* mice and exacerbate glucose intolerance in WT controls

We have previously published that tamoxifen dose-dependent expression of Kras^G12V^ in pancreatic ducts leads to both early and late-stage PanINs and invasive PDAC (10). Similarly, tamoxifen dose determines the extent of acinar Kras^G12V^ recombination in *Acinar^KrasG12V^* mice, with higher doses producing greater recombination, shorter survival, and more severe pancreatic disease progression (**Supplementary Fig. 1A-G**). Consequently, the 1 mg dose was selected for subsequent dietary intervention studies because it induces partial recombination, predominantly PanIN lesions, and extended survival. To determine whether dietary composition influences PDAC development, WT and *Acinar^KrasG12V^*mice, aged 8–10 weeks, were either maintained on SD or randomly assigned to one of three diets: LFD, HFD, or KD. After one month on the assigned diets, *Acinar^KrasG12V^*mice were treated with 1 mg tamoxifen and were monitored for survival and body weight **(Fig. 1A).** KD-fed *Acinar^KrasG12V^* mice had the shortest median survival (26 ± 7 days), significantly lower than the SD-fed (87 ± 29 days, p = 0.02) and LFD-fed mice (57 ± 27 days, p = 0.02). Similarly, HFD feeding reduced survival (35 ± 25 days, p = 0.05) compared with SD **(Fig. 1B)**. At the humane endpoint, KD-fed *Acinar^KrasG12V^* mice exhibited significant weight loss compared with KD-fed WT mice (p = 0.05) **(Fig 1C)**. As expected, KD feeding significantly elevated serum β-hydroxybutyrate (β-OHB) in WT mice compared to SD (p = 0.003), and HFD (p = 0.02) with a similar trend in *Acinar^KrasG12V^*mice (KD vs SD and HFD, p=0.0001). Furthermore, KD-fed *Acinar^KrasG12V^*mice exhibited higher β-OHB than KD-fed WT controls (p = 0.03) **(Fig. 1D)**. In WT mice, KD caused severe glucose intolerance, reflected by a significantly increased area under the glucose-tolerance curve (AUC) compared with SD, LFD, and HFD (p < 0.0001). HFD also impaired glucose tolerance relative to SD and LFD (p < 0.0001). In *Acinar^KrasG12V^* mice, KD reduced glucose tolerance compared to LFD (p = 0.02) and HFD (p = 0.001). When comparing across genotypes, *Acinar^KrasG12V^* mice displayed lower glucose AUC than WT mice under KD (p < 0.0001), and HFD (p < 0.0001) **(Fig.1E)**. The Pancreas to body weight ratio was increased in KD-fed *Acinar^KrasG12V^*compared with WT, whereas the SD-fed *Acinar^KrasG12V^* mice showed reduced pancreas weight compared to WT controls **(Fig. 1F).** No significant differences were observed in serum acetate, amylase or aspartate aminotransferase (AST) levels across diets in either genotype **(Supplementary Fig. S2A-C)**.

**Figure 1.**
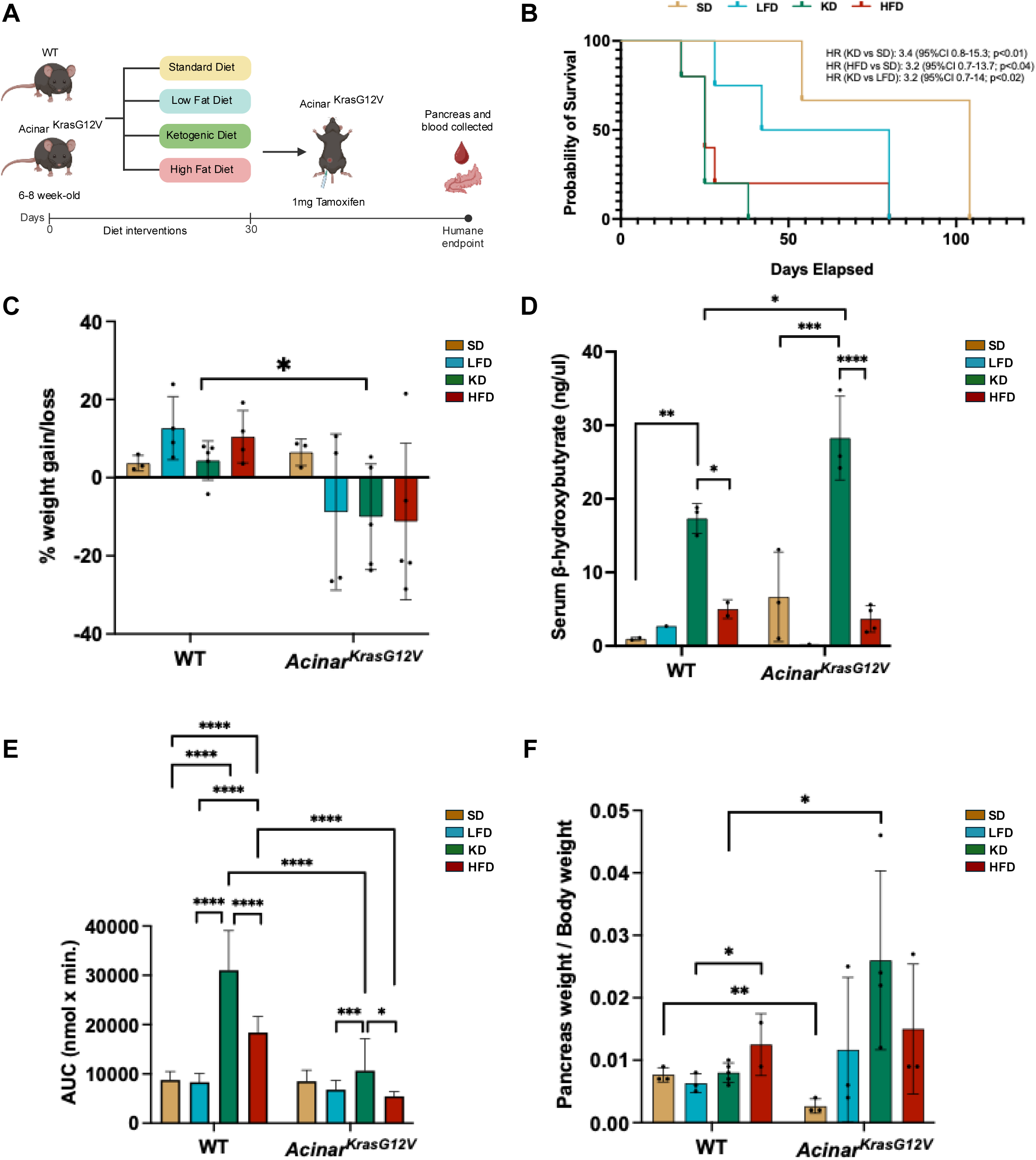
KD- and HFD-fed mice show reduced survival in *Acinar^KrasG12V^* mice and increased glucose intolerance in WT controls. **A)** Experimental design. WT and *Acinar^KrasG12V^* mice (8–10 weeks old) were fed SD, LFD, KD, or HFD for 30 days. *Acinar^KrasG12V^*mice received 1mg of tamoxifen to induce oncogenic Kras activation. Pancreas and serum were collected at the humane endpoint. **B)** Kaplan–Meier survival curves showing probability of survival for *Acinar^KrasG12V^* mice under different dietary interventions. **C)** Percent change in body weight at the endpoint following diet interventions in WT and *Acinar^KrasG12V^* mice. **D)** Serum β-OHB levels at the endpoint in WT and AcinarKrasG12V mice under each diet. **E)** Glucose tolerance test (GTT) area under the curve (AUC) measured 3 weeks after the tamoxifen induction in the WT and *Acinar^KrasG12V^* mice. **F)** Pancreas weight relative to body weight at the end point across dietary groups in WT and *Acinar^KrasG12V^*mice. Statistical analyses were performed using unpaired t-tests (C), one-way ANOVA with Sidak’s test (D), one-way ANOVA with Tukey’s test and two-way ANOVA with Sidak’s test (E), and unpaired t test and one-way ANOVA with Tukey’s test (F). Data are presented as mean ± SEM. *p<0.05, **p<0.01, ***p<0.001, ****p<0.0001.

### KD and HFD promote fibrosis and progression to PDAC in non-obese *Acinar^KrasG12V^* mice

Since KD and HFD-fed *Acinar^KrasG12V^* mice exhibited the worst survival outcomes, we next examined tissue-level changes using histological and immunohistochemical analyses. **Fig. 2A** shows the representative images of ADM, PanIN and PDAC, cytokeratin 19 (CK19), trichrome, CD8, CD39 and CD73 staining in pancreatic tissue sections from *Acinar^KrasG12V^*mice under different diets Pathological evaluation revealed distinct differences in lesion grade and tissue architecture (**Supplemental Table S2)**. The proportion of pancreas replaced by invasive, sarcomatoid-like PDAC was markedly higher in KD-fed *Acinar^KrasG12V^* mice compared to SD-fed mice, which had moderately differentiated neoplasia (p < 0.0001) and LFD-fed mice that presented with well to moderately differentiated PDAC (p = 0.01). HFD-fed *Acinar^KrasG12V^* mice also showed an increase in poorly differentiated PDAC compared to SD (p = 0.0007) **(Fig. 2B)**. These data indicate that both KD and HFD accelerate the development of aggressive and invasive PDAC in *Acinar^KrasG12V^* mice. The percentage of CK19⁺ area, indicating ductal-like differentiation was elevated in LFD, KD and HFD-fed *Acinar^KrasG12V^* mice compared to SD (p = 0.02, p = 0.002, and p = 0.003, respectively **(Fig. 2C)**. Fibrosis, assessed by collagen⁺ area, was markedly increased in KD-fed *Acinar^KrasG12V^* mice, which exhibited the strongest fibrotic response (p = 0.005 vs. SD; p = 0.0002 vs. LFD). Compared to LFD, HFD-fed *Acinar^KrasG12V^*mice also showed significantly higher fibrosis (p = 0.01) **(Fig. 2D)**. Further, there were significantly fewer CD8+ T cells in HFD-fed *Acinar^KrasG12V^*mice compared to SD, indicating HFD may suppress immune responses within the tumor microenvironment. Stromal CD39 expression was significantly higher in KD- and HFD-fed *Acinar^KrasG12V^*mice compared to their respective tumor compartments, whereas CD73 expression showed no significant differences across diets. Together, these findings demonstrate that dietary composition influences the progression of pancreatic lesions, fibrosis, and immune cell landscape in *Acinar^KrasG12V^*mice, with KD and HFD promoting more aggressive disease phenotypes.

**Figure 2.**
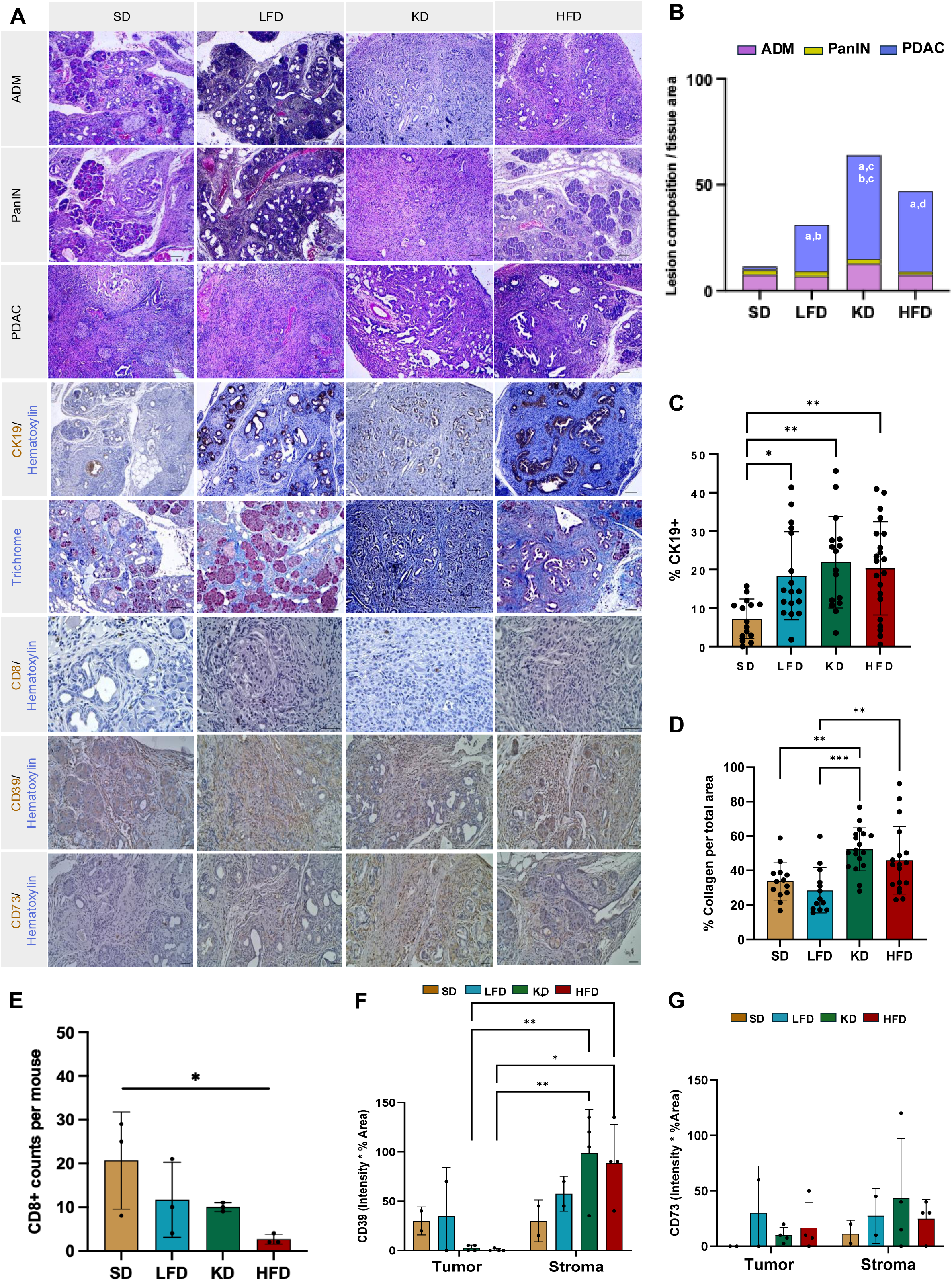
KD and HFD promote fibrosis and progression to PDAC in *Acinar^KrasG12V^* mice. **A)** Representative images of pancreatic tissue from SD, LFD, KD, and HFD-fed *Acinar^KrasG12V^*mice showing ADM, PanIN lesions, PDAC, CK19 immunostaining, trichrome staining, and CD8, CD39, and CD73 expression. Scale bars, 50 µm. **B)** Stacked bar graph depicting the percentage of ADM, PanIN, and PDAC across dietary groups. Quantification of **C)** CK19⁺ epithelial area, **D)** collagen deposition (trichrome⁺ area), and **E)** CD8⁺ T cells counts per mouse in SD, LFD, KD, and HFD-fed *Acinar^KrasG12V^*mice. **F)** CD39 expression (intensity × % area) and **G)** CD73 expression (intensity × % area) quantified separately in tumor and stromal compartments across diets. Statistical analyses were performed using one-way ANOVA with group significance denoted by letter pairs (a = SD, b = LFD, c = KD, d = HFD) (B); one-way ANOVA with Sidak’s test (C-E); and two-way ANOVA with Sidak’s test (F and G). Data are presented as mean ± SEM. *p<0.05, **p<0.01, ***p<0.001, ****p<0.0001.

### High-Fat and Ketogenic Diets Reprogram Tumor Signaling and Systemic Cytokine Profiles

Given that both HFD and KD promoted more aggressive histopathological phenotypes with increased fibrosis and reduced immune cell infiltration, we next sought to determine the molecular signaling pathways underlying these diet-induced effects. We performed RPPA analysis of pancreatic tissues and serum cytokine profiling to assess local and systemic alterations in oncogenic and inflammatory signaling. Volcano plots highlighting significant protein alterations (p < 0.01, |log₂FC| > 0.2) in LFD-, KD-, and HFD-fed WT mice compared with SD-fed controls are shown in **Supplementary Fig. S3A**, and those for *Acinar^KrasG12V^* mice are shown in **Fig.3A**. Compared with SD-fed *Acinar^KrasG12V^* mice, LFD-fed mice exhibited 12 altered proteins (7 downregulated, 5 upregulated), KD-fed mice showed 18 altered proteins (8 downregulated, 10 upregulated), and HFD-fed mice displayed 21 altered proteins (10 downregulated, 11 upregulated). Several proteins were commonly dysregulated in both KD- and HFD-fed groups, including upregulation of Akt, GSK3β, phosphorylated S6 (Ser235/236), phosphorylated AMPKα (Thr172), Paxillin, and YTHDF2, and downregulation of PUMA, STAT5A, c-kit, PHGDH, FASN, and ASNS. Moreover, LFD-fed mice showed upregulation of Akt and phosphorylated CREB (Ser133), and downregulation of PUMA, STAT5A, and c-Kit (**Supplementary Table S3; Fig. 3A)**. To identify biological processes affected by these proteomic changes, KEGG pathway enrichment analysis was performed. The top ten enriched KEGG pathways for LFD-, HFD-, and KD-fed *Acinar^KrasG12V^* mice compared with SD are shown in **Figures 3B–D**, respectively. Across all diet groups, several pathways were commonly upregulated, including EGFR tyrosine kinase inhibitor resistance, chemokine signaling, mTOR signaling, PI3K–Akt signaling, insulin resistance, Rap1 signaling, and VEGF signaling pathways (**Supplementary Table S4; Fig. 3B–D)**.

**Figure 3.**
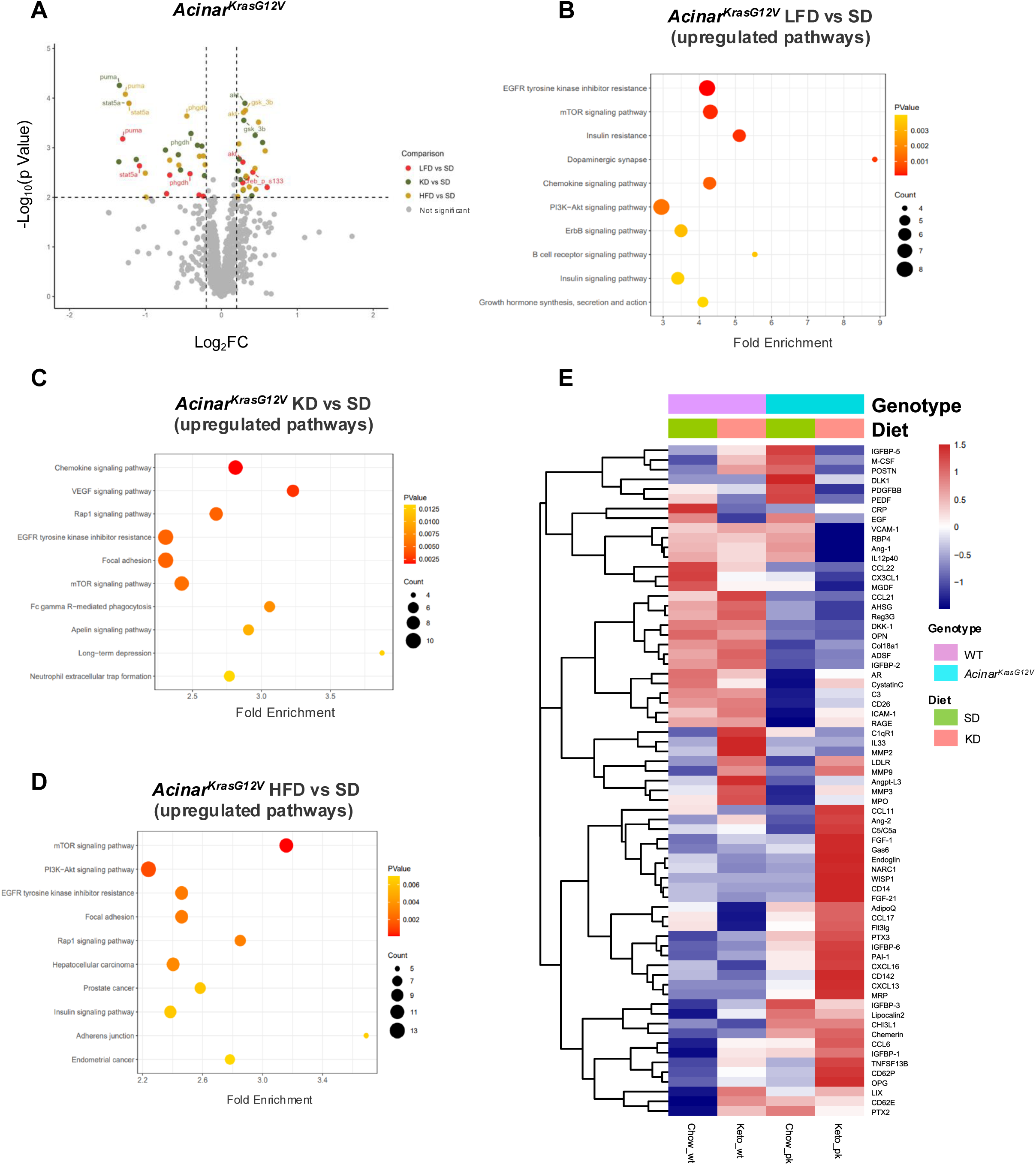
Diet-induced alterations in pancreatic signaling pathways and systemic cytokine profiles in *Acinar^KrasG12V^*mice. **A)** Volcano plot illustrating differential protein expression in *Acinar^KrasG12V^*mice fed LFD, KD, or HFD relative to SD. Proteins were analyzed using the limma package. Significant proteins are highlighted (LFD vs. SD: red; KD vs. SD: green; HFD vs. SD: yellow), and non-significant proteins are shown in grey. The x-axis represents log₂ fold change, and the y-axis shows –log₁₀(*p*) values. Dotted lines indicate p=0.01 and |log₂FC|=0.2 thresholds. KEGG pathway enrichment analysis of upregulated proteins identified in the **(B)** LFD vs. SD, **(C)** KD vs. SD, and **(D)** HFD vs. SD comparisons. Bubble plots display the top enriched pathways ranked by fold enrichment (x-axis). Bubble size corresponds to the number of upregulated proteins contributing to each pathway, and bubble color reflects pathway significance (*p*-value). **(E)** Heatmap depicting serum cytokine and chemokine expression across WT and *Acinar^KrasG12V^*mice fed SD or KD. Cytokines/chemokines are hierarchically clustered by expression pattern. Red indicates higher abundance, while blue indicates lower abundance.

To determine systemic factors underlying accelerated disease progression in KD-fed *Acinar^KrasG12V^* mice, we profiled serum cytokines in SD- and KD-fed WT and *Acinar^KrasG12V^*mice. The heatmap **(Fig. 3E)** revealed a shift in serum cytokine levels in KD-fed groups, with the most profound changes observed in *Acinar^KrasG12V^* mice. The serum levels of Ang-2, CCL6, LDLR, MMP-9, PAI-1, PTX3, TNFSF13B, and OPG were significantly elevated in KD-fed mice and were further increased in KD-fed *Acinar^KrasG12V^* mice compared with their respective SD-fed controls **(Supplementary Fig. S3B)**. Additionally, CCL11, CD14, and FGF-21 were elevated, whereas CX3CL1 and IL-12p40 were reduced in KD-fed *Acinar^KrasG12V^* mice relative to SD-fed counterparts **(Supplementary Fig. S2B)**. Collectively, these findings indicate that the ketogenic diet induces a systemic pro-inflammatory and pro-angiogenic cytokine milieu, coinciding with activation of PI3K–Akt–mTOR and EGFR signaling in the pancreas. Together, these tissue and serum-level alterations provide mechanistic insight into how lipid-rich diet accelerates pancreatic tumor progression in *Acinar^KrasG12V^*mice.

## DISCUSSION

Using an inducible acinar-specific Kras model, we found that pre-tumor dietary exposure profoundly shaped pancreatic disease trajectory. Both the KD and HFD accelerated disease progression, increased invasive PDAC, fibrosis, and reduced survival compared with SD and LFD. Previous studies have shown that acinar cells possess plasticity in response to inflammation and Kras^G12D^ mutations (11–14), and our findings extend this by demonstrating how macronutrient balance before oncogenic activation conditions the tissue’s response to KRAS signaling, with lipid-rich diets favoring aggressive neoplasia.

The detrimental impact of HFD aligns with prior work showing that obesogenic high-fat feeding promotes acinar injury, ADM, PanIN formation, and PDAC progression in Kras^G12D^ mice (1, 2). Our results also support emerging evidence that KD can accelerate PDAC in lean Kras-mutant models despite having opposite effects in obesity-associated settings, highlighting metabolic state as a determinant of KD’s impact (7). KD-fed WT and *Acinar^KrasG12V^* mice showed marked glucose intolerance, consistent with reports in non-obese rodents, and insulin resistance may enhance PI3K–Akt–mTOR signaling that cooperates with KRAS during early neoplastic development.

This context dependency suggests that metabolic state rather than KD itself determines whether very low–carbohydrate, high-fat intake is protective or harmful. In our study, KD-fed WT and *Acinar^KrasG12V^*mice developed significant glucose intolerance, a phenotype also reported in non-obese rodents fed long-term KD (15). Given that insulin resistance can amplify PI3K–Akt–mTOR signaling, and that this pathway cooperates with KRAS to drive early neoplastic progression, diet-induced metabolic stress likely contributed to the enhanced disease in KD-fed mice (16, 17).

Although LFD was less detrimental than KD or HFD, our findings parallel those of previous work demonstrating that carbohydrate-rich diets can also promote early pancreatic lesions (3). The LFD used in our study (77% carbohydrate) is compositionally similar to the high-carbohydrate diet evaluated by Zhu et al., who reported shortened survival and increased ADM, inflammation, and PanIN development in Kras^G12D^ mice (18). Together, our results support the concept that both excessive dietary fat and high proportions of rapidly metabolized carbohydrates can worsen early pancreatic neoplasia, although lipid-rich diets exerted the strongest effects.

KD and HFD increased collagen deposition and stromal CD39 expression, implicating adenosine-mediated immune suppression in accelerated disease (19) (20). KD-associated increases in β-HB may further enhance invasiveness by stabilizing the EMT regulator Snail (21). Systemically, KD elevated pro-inflammatory and pro-angiogenic cytokines that support matrix remodeling, invasion and systemic pro-inflammatory and pro-angiogenic cytokine profiles including elevated Ang-2, MMP-9, PAI-1, PTX3, and TNFSF13B (22–24). Overall, these findings demonstrate that diet strongly shapes early KRAS-driven neoplastic evolution and warrant caution regarding high-fat or ketogenic patterns in individuals at elevated pancreatic cancer risk.

## Methods

### Animal Experiments

All animal experiments were conducted in accordance with the guidelines approved by the Institutional Animal Care and Use Committee at the University of Texas Health Science Center at Houston and the University of Nebraska Medical Center *Pft1aCre^ERT2^ Kras^G12V^* (*Acinar^KrasG12V^)* mice and wild-type C57BL/6J mice were purchased from the Jackson Laboratory. Mice were maintained on a 12-hour light/dark cycle with ad libitum access to a standard chow diet (SD) until dietary intervention.

### Tamoxifen dosing in *Acinar^KrasG12V^* mice

To establish dosing conditions for oncogenic Kras activation, *Acinar^KrasG12V^*mice (8-10 weeks old) received intraperitoneal injections of tamoxifen at doses of 1 mg, 5 mg or 10 mg. Based on the dose administered, animals were designated as Kras^Low^ (1 mg), Kras^Mod^ (5 mg), or Kras^High^ (10 mg). Following treatment, mice were monitored for survival, and pancreatic tissues were collected for assessment of acinar Kras recombination, lesion development, and collagen deposition.

### Diet interventions

At 8 to 10 weeks of age, the mice were either maintained on the SD or randomly assigned to one of three experimental diets: KD (TD.160153, Teklad Custom Diet; 90% energy from fat), HFD (TD.160239, Teklad Custom Diet; 75% energy from fat, with a fat profile modified to better resemble the KD), or KD Control Diet (LFD) (TD.150345, Teklad Custom Diet; 13% energy from fat). In total, the study included 3 WT and 3 *Acinar^KrasG12V^* mice on SD, 4 WT and 4 *Acinar^KrasG12V^* mice on the LFD, 4 WT and 5 *Acinar^KrasG12V^* mice on the HFD, and 5 WT and 5 *Acinar^KrasG12V^* mice on the KD. The calorie content and exact composition of each diet are provided in Supplemental Table S1. Mice were on assigned diet for one month, after which *Acinar^KrasG12V^*mice received a 1 mg intraperitoneal dose of tamoxifen to induce the oncogenic Kras mutation. Body weight was measured weekly to monitor the overall health and weight changes of the animals under different dietary conditions. Mice were fed ad libitum to explore how diet may modulate the earliest stages of pancreatic neoplasia; we performed a controlled dietary intervention in mice and assessed PanIN prevalence after one month of exposure to distinct dietary regimens. This approach allowed us to examine whether short-term dietary modification can influence PanIN development or regression, thereby providing insight into how nutritional factors shape the earliest steps in pancreatic tumorigenesis. Mice remained on their respective diets until reaching the humane endpoint.

### Glucose tolerance test (GTT)

GTT was carried out 10 days after tamoxifen administration. Prior to the test, mice were subjected to a 6-hour fasting period. The glucose dose was calculated based on body weight after fasting (2.0 g of glucose per kilogram of body weight). Blood samples were obtained from the tail vein before glucose injection (baseline) and at 15, 30, 60, and 120 minutes afterward. Blood glucose concentrations were determined using a Contour Next One Blood Glucose Monitoring System.

### H&E staining

Formalin-fixed, paraffin-embedded tissues were sectioned and mounted on charged glass slides. Slides were deparaffinized in Histoclear, rehydrated through a graded ethanol to water, and stained with hematoxylin. Slides were then counterstained with eosin, dehydrated in ethanol, cleared in Histoclear, and mounted.

### IHC

Slides were baked at 60°C, deparaffinized in Histoclear, rehydrated through graded ethanol to PBS and subjected to heat-mediated antigen retrieval using either citrate buffer (pH 6.0) or Tris-EDTA buffer (pH 9.0). Sections were blocked for 1 hour at room temperature in 10% FBS prepared in PBST and then incubated overnight at 4°C with primary antibodies diluted in blocking solution. The following day, slides were washed in PBS and incubated for 30 minutes at room temperature with biotinylated secondary antibodies (1:500, Vector Laboratories). Signal detection was performed using the Vectastain Elite ABC kit (Vector Laboratories) followed by the DAB substrate (Vector Laboratories). Sections were counterstained with hematoxylin, dehydrated through graded ethanol, cleared in Histoclear, and mounted.

### Trichrome staining

Connective tissue was visualized using the Abcam Trichrome Stain Kit (ab150686) according to the manufacturer’s protocol. Briefly, slides were deparaffinized in Histoclear, rehydrated through graded ethanol to distilled water, and incubated in preheated Bouin’s solution (60°C) for 60 minutes. After rinsing in tap water, sections were stained sequentially with Weigert’s iron hematoxylin (5 minutes), Biebrich Scarlet/acid fuchsin (15 minutes), phosphomolybdic–phosphotungstic acid (10–12 minutes), and aniline blue (10–20 minutes). Sections were then treated with acetic acid for 5 minutes, dehydrated through graded ethanol, cleared in Histoclear, and mounted.

### Image J analysis

Quantification of IHC staining was performed using ImageJ software (https://imagej.net/software/fiji/).

5–10 representative fields were selected depending on tissue size. Positive staining was identified using the color threshold tool, which normalized signal across tissues. For analyses of ADM, PanIN or PDAC in H&E-stained sections, the freehand tool was similarly used to outline tissue regions of interest, and their relative areas were measured.

### Reverse Phase Protein Assay (RPPA)

Small pieces of frozen pancreatic tissue were homogenized in ice-cold lysis buffer (1% Triton X-100, 50mM HEPES pH 7.4, 150mM NaCl, 1.5mM MgCl2, 1mM EGTA, 100mM NaF, 10mM Na pyrophosphate, 1mM Na_3_VO_4_, 10% glycerol) supplemented with protease and phosphatase inhibitors. Lysates were centrifuged at 14,000 rpm for 10 minutes at 4°C, and the supernatant was collected. Protein concentration was measured by BCA assay and adjusted to 1.5 µg/µl with lysis buffer. Samples were combined with 4× SDS sample buffer (40% glycerol, 8% SDS, 0.25 M Tris-HCl, pH 6.8) containing β-mercaptoethanol at a 3:1 ratio (sample:buffer), boiled for 5 minutes, and stored at –80°C until analysis.

RPPA analysis was performed at the Functional Proteomics Core Facility at The University of Texas MD Anderson Cancer Center. Protein lysates were serially diluted in five two-fold steps and arrayed onto nitrocellulose-coated slides (Grace Bio-Labs) using a Quanterix 2470 Arrayer, generating 5,808 spots per slide. Each slide included experimental samples, standard lysates, and positive/negative controls. Each slide was probed with a validated primary antibody plus a biotin-conjugated secondary antibody. Signal was amplified with the Agilent GenPoint system and visualized by DAB colorimetric reaction. Slides were scanned using a Huron TissueScope, and spot intensities were quantified with Array-Pro Analyzer software (Media Cybernetics).

Relative protein expression was determined using RPPA SPACE software (MD Anderson Department of Bioinformatics and Computational Biology) through logistic “supercurve” fitting of the serial dilution data. A single fitted curve was generated for each antibody across all samples, with dilution step as the independent variable and signal intensity as the response. Protein levels were normalized for loading variation by bidirectional median centering (across samples for each antibody and across antibodies for each sample). Replicates-based normalization (RBN) using control samples was further applied to correct for inter-set variation, allowing direct comparison across different RPPA batches (25).

Differential protein expression across diet groups in WT and *Acinar^KrasG12V^*mice was assessed using the limma package in R (version 4.5.2). Multiple hypothesis testing correction was performed using the Benjamini–Hochberg method to control the false discovery rate (FDR). Volcano plots were generated using a significance threshold of p < 0.01 and |log₂FC|=0.2 . For pathway enrichment analysis, proteins were filtered using p < 0.05 and |log2FC| > 0.10. Volcano plots and pathway analysis was performed using ggplot2 and clusterProfiler and org.Mm.eg.db annotation package (Mus musculus).

### Serum cytokine array

Serum cytokine and chemokine profiles were analyzed using the Proteome Profiler Mouse XL Cytokine Array kit (ARY028; R&D Systems, Minneapolis, MN, USA), which detects the relative expression of 111 cytokines/chemokines simultaneously. The assay was performed according to the manufacturer’s protocol, and chemiluminescent signal intensity was used to determine relative expression levels. For this analysis, one serum sample from each group WT SD, *Acinar^KrasG12V^* SD, WT KD, and *Acinar^KrasG12V^*KD was included. Imaging was performed on a Bio-Rad ChemiDoc instrument. Heatmap of serum cytokine expression were generated using the pheatmap package in R. Statistical comparisons were performed using two-way ANOVA, followed by Tukey’s multiple comparisons test.

### Serum Acetate, β-OHB, Amylase and AST levels

Serum acetate levels were quantified using the Acetate Colorimetric Assay Kit (Millipore Sigma, catalog no. MAK086). Serum β-hydroxybutyrate (β-OHB) levels were measured using the β-Hydroxybutyrate Assay Kit (Millipore Sigma, catalog no. MAK041). Serum amylase levels were quantified using the Pointe Liquid Amylase (CNPG3) Reagent Set (MedTest DX). Aspartate aminotransferase (AST) levels were measured using the Teco Diagnostics AST Assay Kit (catalog no. A559-150). All assays were performed according to the manufacturers’ instructions.

## Acknowledgements

Research reported in this publication was supported by the National Cancer Institute of the National Institutes of Health under award number P30 CA036727 (UNMC CCSG Cancer Center Grant). The content is solely the responsibility of the authors and does not necessarily represent the official views of the National Institutes of Health. Data were generated in part through the use of the Functional Proteomics RPPA Core (RPPA), which receives partial support from the National Cancer Institute under grant P30CA016672 to MD Anderson Cancer Center. The research reported in this publication was not directly funded through the grant P30CA016672 to MD Anderson Cancer Center and is not within the scope of such grant. JMB-L received funding from (R01CA277161-01A1, R21CA249924) and DOD (HT94252410921). FM received support from the National Cancer Institute (1R37CA237384, R01 CA282786), Cancer Prevention Research Institute of Texas (RP200173), and DOD (HT94252310665).

