## Supplemental Figures for "Dietary Fat Content Influences PanIN Progression and Pancreatic Cancer Development in Mice"

### Supplementary Figure S1

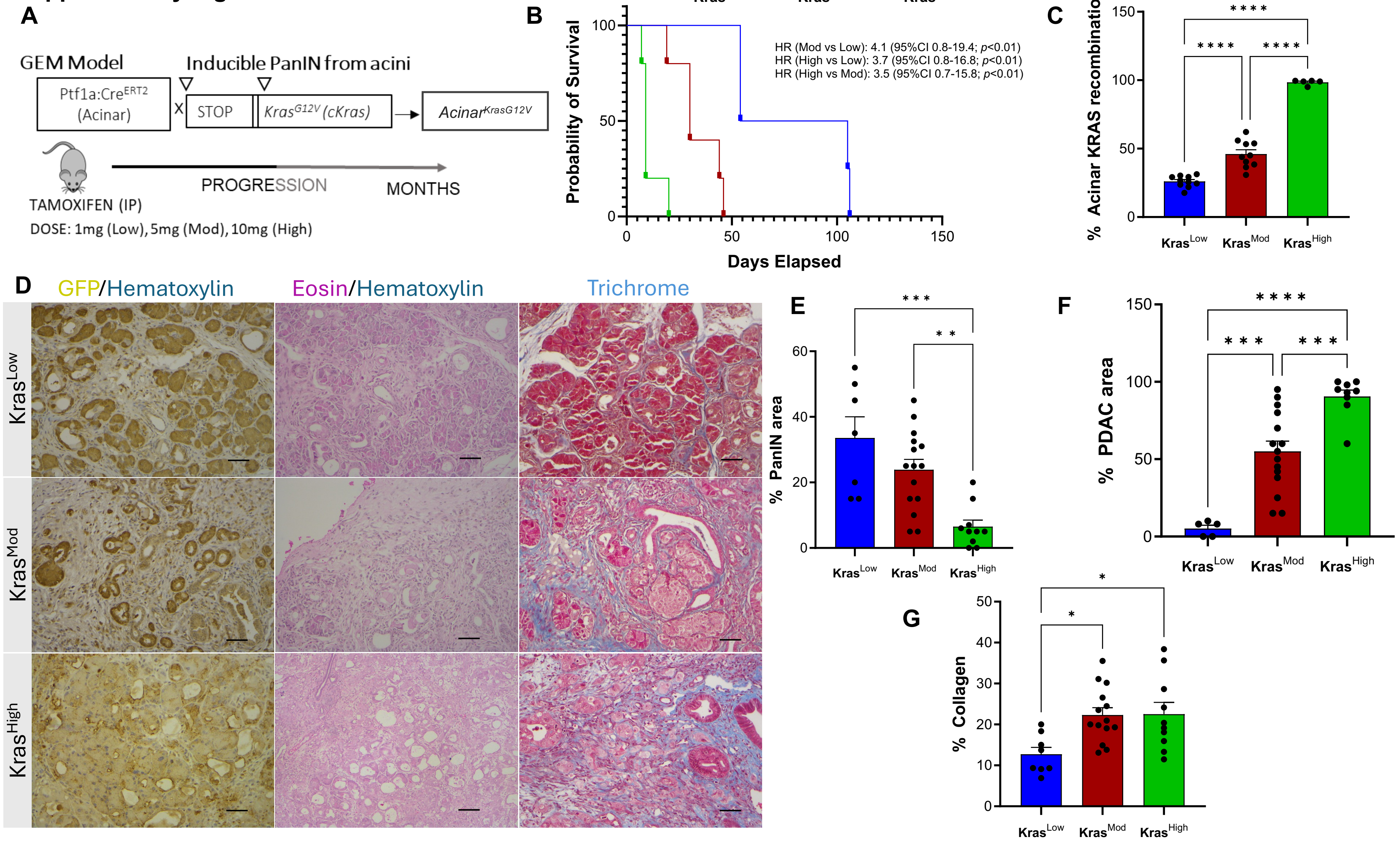


**Figure S1.** **Dose-dependent effects of tamoxifen on Kras recombination, survival, and pancreatic pathology in *Acinar*$^{KrasG12V}$ mice. A)** Schematic of the experimental plan showing the effect of varying tamoxifen doses (1 mg, Kras$^{Low}$; 5 mg, Kras$^{Mod}$; 10 mg, Kras$^{High}$) on Kras recombination efficiency in *Acinar*$^{KrasG12V}$ mice. **B)** Kaplan–Meier survival curve showing survival of *Acinar*$^{KrasG12V}$ mice treated with different tamoxifen doses. **C)** Percentage of acinar KRAS recombination in Kras$^{Low}$, Kras$^{Mod}$ and Kras$^{High}$ mice, as assessed by GFP loss. **D)** Representative images showing H&E and Trichrome staining of pancreatic tissue from Kras$^{Low}$, Kras$^{Mod}$ and Kras$^{High}$ mice. Scale bars are 50μM. Percentage of **E)** PanIN lesions **F)** PDAC and **G)** Collagen deposition in Kras$^{Low}$, Kras$^{Mod}$ and Kras$^{High}$ mice. Statistical significance was determined by one way ANOVA with Tukey's multiple comparison test. *p<0.05, **p<0.01, ***p<0.001, ****p<0.0001. n= 4 mice in Kras$^{Low}$ group and n = 5 mice in Kras$^{Mod}$ and Kras$^{High}$ group.

### Supplementary Figure S2

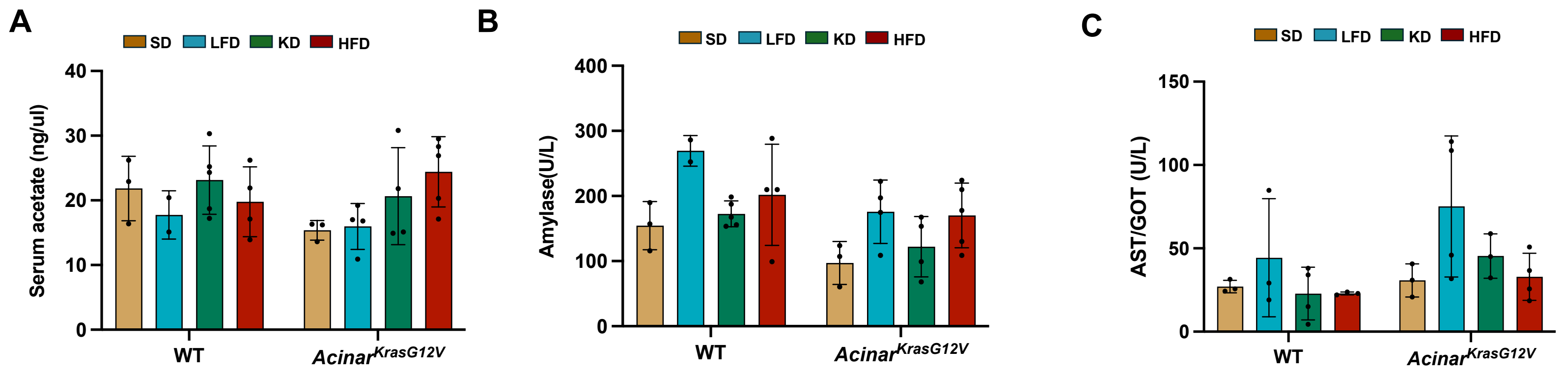


**<u>Figure S2.</u> Serum acetate and amylase levels in WT and *Acinar*$^{KrasG12V}$ mice across dietary groups**. **A)** Serum acetate levels at the endpoint in WT and *Acinar*$^{KrasG12V}$ mice under each diet. **B)** Serum amylase levels at the endpoint in WT and *Acinar*$^{KrasG12V}$ mice under each diet. **C)** Serum AST levels at the endpoint in WT and *Acinar*$^{KrasG12V}$ mice under each diet.

### Supplementary Figure S3

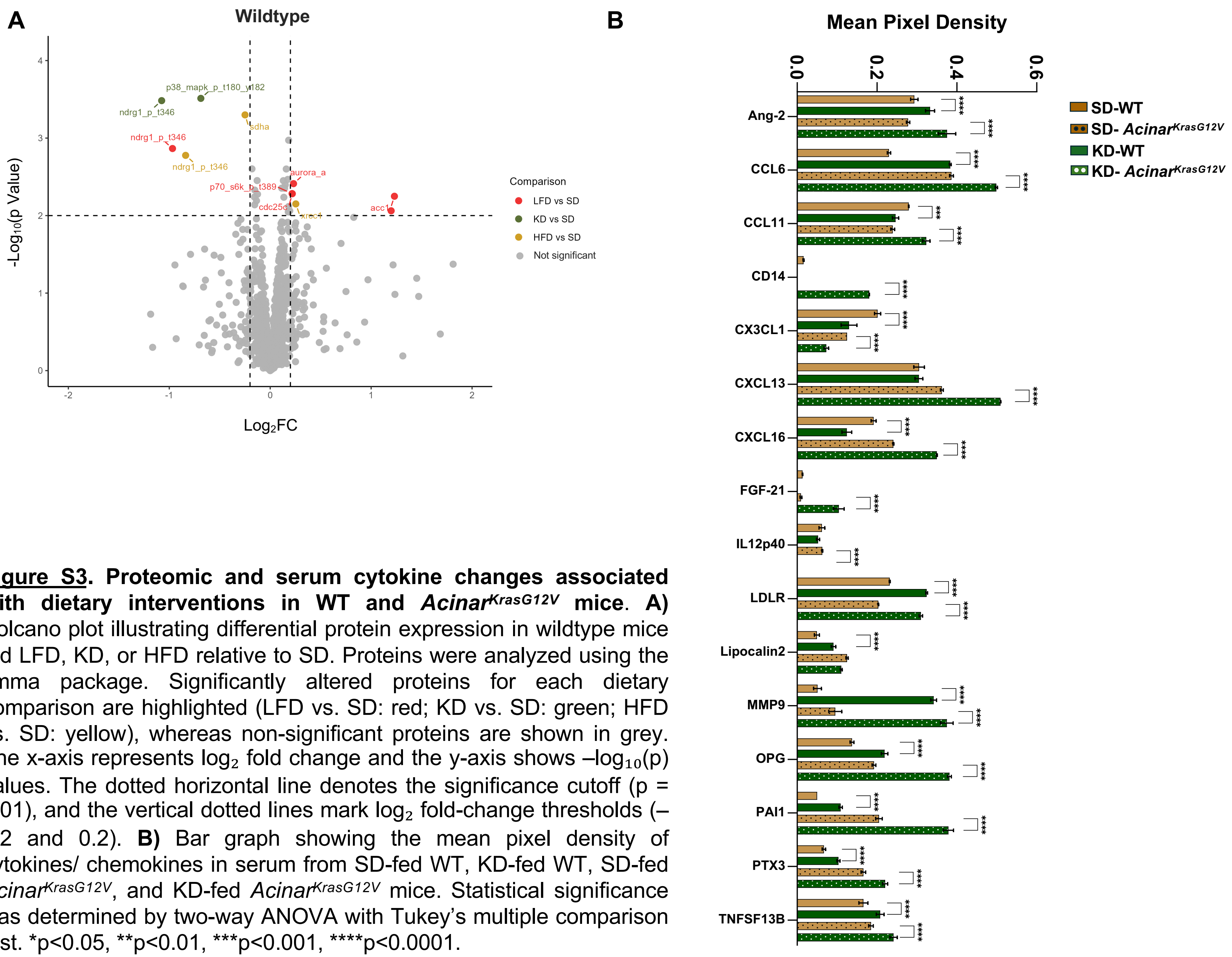


__Figure S3__. **Proteomic and serum cytokine changes associated with dietary interventions in WT and *Acinar*$^{KrasG12V}$ mice. A)** Volcano plot illustrating differential protein expression in wildtype mice fed LFD, KD, or HFD relative to SD. Proteins were analyzed using the limma package. Significantly altered proteins for each dietary comparison are highlighted (LFD vs. SD: red; KD vs. SD: green; HFD vs. SD: yellow), whereas non-significant proteins are shown in grey. The x-axis represents $\log_2$ fold change and the y-axis shows $-\log_{10}(p)$ values. The dotted horizontal line denotes the significance cutoff ($p = 0.01$), and the vertical dotted lines mark $\log_2$ fold-change thresholds (–0.2 and 0.2). **B)** Bar graph showing the mean pixel density of cytokines/ chemokines in serum from SD-fed WT, KD-fed WT, SD-fed *Acinar*$^{KrasG12V}$, and KD-fed *Acinar*$^{KrasG12V}$ mice. Statistical significance was determined by two-way ANOVA with Tukey's multiple comparison test. *p<0.05, **p<0.01, ***p<0.001, ****p<0.0001.
