## Supplemental Table 1 for "Dietary Fat Content Influences PanIN Progression and Pancreatic Cancer Development in Mice"

|  | **Ketogenic 93M Diet (KD)**  **TD.160153** | **High Fat Diet**  **(HFD)**  **TD.160239** | **KD Control or**  **Low Fat Diet (LFD)**  **TD.150345** |
| --- | --- | --- | --- |
| **Ingredients** | **g/kg** | **g/kg** | **g/kg** |
| Casein | 180 | 160 | 100 |
| DL-Methionine | 2.88 | 2.5 | 1.6 |
| Vegetable Shortening, hydrogenated (Crisco) | 440 | 314.0 | 25 |
| Cocoa Butter | 150 | 107.0 | 0 |
| Corn Oil | 85 | 60.715 | 25 |
| Cellulose | 59.1884 | 49.3168 | 34.918 |
| Vitamin Mix, AIN-93-VX w/ Cellulose (110068) | 27 | 24 | 15 |
| Thiamin (81%) | 0.018 | 0.016 | 0.01 |
| Vitamin K1, phylloquinone | 0.0036 | 0.0032 | 0.002 |
| Choline Bitartrate | 4.5 | 3.85 | 2.5 |
| Mineral Mix, w/o CA & P (98057) | 24.1 | 20.887 | 13.39 |
| Calcium Phosphate, dibasic | 17.64 | 15 | 9.8 |
| Calcium Carbonate | 9.54 | 8.1 | 5.25 |
| TBHQ, antioxidant | 0.13 | 0.112 | 0.07 |
| Corn Starch | 0 | 158 | 512.46 |
| Sucrose | 0 | 30 | 100 |
| Maltodextrin | 0 | 46.5 | 155 |
| **Macronutrient energy** | **% kcal** | **% kcal** | **% kcal** |
| Protein | 9.5 | 9.8 | 9.7 |
| Carbohydrate | 0 | 15.0 | 77.7 |
| Fat | 90.5 | 75.2 | 12.6 |

**Supplementary Table 1:** Diet compositions. The table lists ingredient amounts (g/kg) and the relative contributions of protein, carbohydrate and fat to the total calories of each diet.

Diets used in this study were from Teklad Custom Diet
