## Supplemental Table 2 for "Dietary Fat Content Influences PanIN Progression and Pancreatic Cancer Development in Mice"

**Supplementary Table 2:** Pathological evaluation of pancreatic tissue sections from *Acinar^KrasG12V^* mice under different diets.

| **Diet** | **Pathological Description** |
| --- | --- |
| SD | Predominantly normal acinar architecture with residual acini visible; minimal evidence of malignancy; mild to moderate inflammatory infiltrate; moderately differentiated lesions; islet cell hyperplasia. |
| LFD | Areas of malignancy involving acinar regions; mild inflammation; overall well to moderately differentiated lesions. |
| KD | Presence of sarcomatoid transformation, mild to moderate inflammation, neutrophils infiltration leading to cell death and release of DAMPs; increased stemness observed. |
| HFD | Marked malignancy with poorly differentiated lesions; minimal inflammation observed. |
